# Reconstructing Sample-Specific Networks using LIONESS

**DOI:** 10.1101/2021.09.27.461954

**Authors:** Marieke L. Kuijjer, Kimberly Glass

## Abstract

We recently developed LIONESS, a method to estimate sample-specific networks based on the output of an aggregate network reconstruction approach. In this manuscript, we describe how to apply LIONESS to different network reconstruction algorithms and data types. We highlight how decisions related to data preprocessing may affect the output networks, discuss expected outcomes, and give examples of how to analyze and compare single sample networks.

## Introduction

Network reconstruction methods, especially those based on transcriptomic data, generally estimate a single network for a given set of input samples by applying an aggregate statistic to the data. Although aggregate networks have had a profound impact on our understanding of biological systems, the population-level networks estimated by most reconstruction approaches cannot, by definition, provide information on the specific network processes that are active in each individual person.

We recently developed a method, called LIONESS (Linear Interpolation to Obtain Network Estimates for Single Samples), that addresses this gap by using a leave-one-out approach to reverse engineer sample-specific networks (Kuijjer et al., 2019a). The basic idea of LIONESS is to estimate a sample-specific network by incorporating information regarding both the similarities and the differences between two networks—one modeled with and one modeled without the sample of interest. LIONESS determines how the removal of this single sample perturbs the network estimated on all samples. It then combines this information with the network modeled on all samples except the sample of interest. This combination makes the LIONESS approach distinct from methods that only estimate edges that are specific to a sample by calculating perturbations (Liu et al., 2016). LIONESS models *all* of the edges in each single-sample network, including those that are commonly shared across all samples.

The single-sample networks predicted by LIONESS provide a way to unite (i) the extensive literature and methodologies for estimating complex network relationships using genomic data with (ii) statistical analysis techniques that use sample-level information to model heterogeneity and compare phenotypic groups. Applications of LIONESS have included analyzing the yeast cell cycle (Kuijjer et al., 2019a), studying biological processes in lymphobastoid cell lines (Kuijjer et al., 2019a), investigating the relationship between the host transcriptome and nasal microbiome in infants with respiratory syncytial virus infection (Sonawane et al., 2019), investigating microbiome co-occurrence in respiratory infections (Mac Aogáin et al., 2021), identifying regulatory processes associated with brain cancer survival (Lopes-Ramos et al., 2021) and with sexual dimorphism in colorectal cancer chemotherapy response (Lopes-Ramos et al., 2018), characterizing sex differences in twenty-nine tissues from the Genotype-Tissue Expression Project (Lopes-Ramos et al., 2020), and identifying tissue-specific regulatory processes in maize (Fagny et al., 2020). LIONESS has also been integrated into a method to identify cancer driver genes (Pham et al., 2021). In many of these applications, LIONESS networks have been analyzed with linear models to statistically evaluate changes in networks in the context of relevant biological and phenotypic variables, as well as potential confounders. This is a significant strength of the downstream use of the single-sample networks predicted using LIONESS.

Importantly, LIONESS is not itself a network reconstruction approach. Rather, it takes the output of an existing reconstruction approach and estimates single-sample versions of those networks in addition to the aggregate model. This means that LIONESS can be applied to many different types of networks, such as co-expression, co-abundance, and regulatory networks. However, this also means that the single-sample networks estimated using LIONESS will be greatly influenced by the choice of the underlying reconstruction approach. Similarly, the single-sample networks estimated by LIONESS depend on the background dataset, just as any reconstructed aggregate network does.

In this manuscript, we demonstrate the application of LIONESS to two data sets—one in human and one in yeast—using two different aggregate network reconstruction approaches—Pearson correlation and PANDA (Passing Attributes between Networks for Data Assimilation, Glass et al., 2013). These approaches make use of transcriptomics data to estimate a gene co-expression (Pearson) or regulatory network (PANDA). Many of the caveats we discuss in this manuscript relate to important decisions that are made when using these aggregate approaches and processing their corresponding input data, since it is critical to understand how these impact the single-sample network models predicted using LIONESS. However, we note that the general framework we lay out in this paper for running LIONESS and evaluating its single-sample network predictions is applicable when using LIONESS together with other aggregate approaches and/or (omics) data types. The code we use for the analyses presented in this paper is available at https://github.com/kuijjerlab/LIONESS_StarProtocols.

### General Aspects of LIONESS

#### Mathematical Framework

LIONESS works by iteratively leaving out a single sample, inferring two aggregate networks (one with and one without the sample) and then using these two networks to estimate a network for the missing sample:

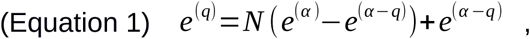

where is *e*^(q)^ the adjacency matrix for the network for sample *q, e*^(*α*)^ is the adjacency matrix for a network reconstructed using all samples, and *e*^(*α −q*)^ is the adjacency matrix for a network reconstructed using all samples except for sample *q*. Based on this flexible framework, LIONESS can extract single-sample networks for the vast majority of aggregate network reconstruction methods, including simple statistical approaches such as Pearson or Spearman correlation (Kuijjer et al., 2019b; Sonawane et al., 2019; Mac Aogáin et al., 2021) and mutual information (Kuijjer et al., 2019a).

#### Procedure and Pseudocode

The LIONESS approach has been implemented in multiple programming languages. Because LIONESS is a wrapper that is applied to a pair of aggregate networks, these implementations occur within software packages that include other network reconstruction algorithms. For example, implementations of LIONESS that can be applied to Pearson correlation networks are available in R (Kuijjer et al., 2019b) and Python (van IJzendoorn et al., 2016). Implementations of LIONESS that can be applied to PANDA are available in R (netzoo.github.io/), MATLAB (Glass et al., 2015), and Python (van IJzendoorn et al., 2016). However, we would like to emphasize that LIONESS is not limited to these languages or applications to these reconstruction approaches.

The LIONESS equation is derived for each edge independently. However, network reconstruction approaches, especially those that use transcriptomics data, generally estimate values for all possible edges in a network. In this case, the LIONESS procedure can be described in the following four steps (see also Figure 1A):

**Figure 1.**
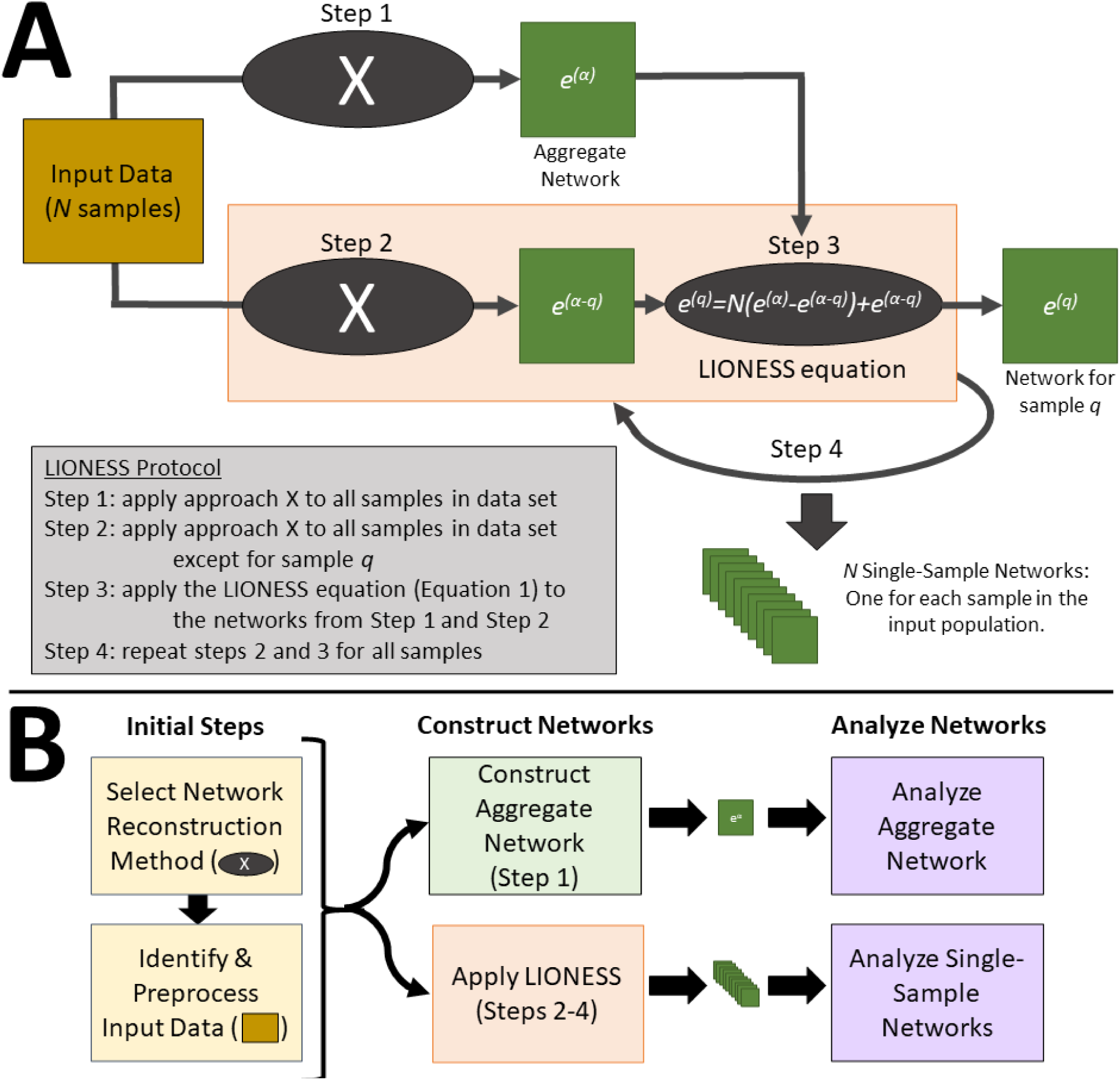
(A) An overview of the LIONESS procedure. LIONESS starts with Input Data for *N* samples, applies a network reconstruction approach (X) determined by the user to estimate two aggregate networks, one based on all the input samples (*e*^(*α*)^) and one based on all the input samples except for sample *q* (*e* ^(*α − q*)^). It then repeats this for each sample in the Input Data to generate *N* total single-sample networks. (B) An overview of how to apply LIONESS in the context of single-sample network analysis. Before applying LIONESS, the user must first specify a network reconstruction approach (X) to use in the procedure and identify/preprocess all the Input Data required for that approach. Next, LIONESS can be used to estimate both the aggregate network (*e* ^(*α*)^) representing information from all samples, as well as *N* single-sample networks. Finally, these networks can be analyzed using both network metrics and statistical techniques.

Step 1: apply approach X to estimate a network using all of the samples in a data set

Step 2: apply approach X to estimate a network using all of the samples in a data set except for sample q

Step 3: apply the LIONESS equation (Equation 1) to the networks from Step 1 and Step 2 (optional): print the single-sample network to a file

Step 4: repeat steps 2 and 3 for all samples

The output of this four step procedure will be a set of single-sample networks, one for each sample in the input dataset. Note that this procedure assumes that the outputs of steps one and two include real values for every possible edge in the network.

Based on this procedure, LIONESS can be implemented in any programming language using the following pseudocode:

1. NetA=NetworkReconstructionApproach(AllSamples);
2. for (Q = 1 to NumSamples)
3. NetAminusQ = NetworkReconstructionApproach(AllSamples except Q)
4. NetQ = NumSamples * (NetA-NetAminusQ) + NetAminusQ

In the sections below, we describe several initial steps that are required prior to running LIONESS, considerations to take into account when constructing single-sample networks using LIONESS, as well as approaches for single-sample network analyses (Figure 1B).

### Choosing a network reconstruction approach and identifying the required data

As noted above, LIONESS is not a standalone procedure, but only works when applied in synergy with an existing network reconstruction algorithm. Therefore, as an initial step prior to running LIONESS, it is necessary to choose a network reconstruction algorithm and to understand its strengths, weaknesses, limitations, and requirements—LIONESS will inherit and amplify these features. For example, an approach that is computationally expensive may require additional preprocessing, memory, or data storage requirements to allow it to scale effectively with LIONESS. In addition, any mathematical features of the aggregate network—such as embedded patterns or negative relationships—will propagate into the LIONESS-estimated networks and will need to be properly modeled in the downstream analyses. LIONESS is most effective when it is applied to an algorithm that estimates a weight for every possible edge.

Different “classes” of network reconstruction approaches exist, including linear and nonlinear methods, methods that model networks on single-omics data, and approaches that integrate multiple data types (Yan et al., 2017). Below, we will demonstrate the application of LIONESS on two of these classes of approaches. We will use Pearson correlation as an example of a linear, single-omics approach, and PANDA as an example of an integrative, non-linear approach. We note that many of the principles we describe in the discussion on Pearson correlation networks can be applied to other single-omics symmetric network estimations, such as Weighted Gene Co-expression Network Analysis (Langfelder and Horvath, 2008), Mutual Information (MI), Context Likelihood of Relatedness (Faith et al., 2007), or Algorithm for the Reconstruction of Accurate Cellular Networks (Margolin et al., 2006), although each specific method will also have additional caveats that will need to be explored (see the *Expected outcomes* section). Similarly, many of the issues that we discuss below in the context of applying LIONESS to PANDA networks are related to the initial data processing and will be necessary to consider when applying LIONESS to other complex methods.

#### Tip

Always familiarize yourself with the selected network reconstruction approach before using it together with LIONESS. Before applying LIONESS to generate single-sample networks, it is useful to first construct and characterize the aggregate network reconstructed using all samples.

#### Required data types

Before running LIONESS, you will first need to collect the correct input data types for the selected network reconstruction algorithm. Below, we describe this process for two approaches —modeling co-expression networks with Pearson correlation and modeling gene regulatory networks with PANDA.

##### Pearson correlation

To model co-expression networks using Pearson correlation, gene expression data—for example obtained from RNA-Seq or microarray experiments—is the only required input.

##### PANDA

PANDA models gene regulatory networks by integrating information on putative regulatory interactions with co-expression patterns (which are modeled using Pearson correlation). PANDA therefore requires both gene expression data, as well as prior information on putative regulatory interactions (also called a “prior regulatory network”) as input. The latter can, for example, be derived from a transcription factor (TF) motif scan, where the DNA-binding motifs of a TF are scanned for within promoter regions to identify potential TF binding sites. In addition, PANDA can integrate protein-protein interactions (PPIs) between the regulators (e.g. TFs). Although this data type is not required, in this paper we choose to include this data type in modeling single-sample networks based on PANDA.

#### Minimum sample size

The minimum sample size required to run LIONESS depends on the underlying network reconstruction approach. For both Pearson correlation and PANDA, the absolute minimum number of samples would be 3+1, since the minimum number of samples to calculate the Pearson correlation coefficient is three, and one sample will be iteratively removed when running LIONESS. However, it is generally recommended to model networks on enough samples to make a reliable aggregate network model. We previously showed that including 20 samples results in a reasonable network similarity score for LIONESS networks built on Pearson or PANDA (Kuijjer et al., 2019a). This reproducibility increased when including more samples.

##### Tip

Should you have a dataset with very few samples, it can be combined with external data from the same or a similar condition (e.g. samples derived from the same disease or cell type) to increase power for building LIONESS networks. In this case, the data should be properly merged and batch-corrected (Kuijjer et al., 2019a) to ensure that edges in the aggregate network are not technical artifacts (see *Preprocessing* section below).

#### What samples to include

LIONESS uses a “background” set of samples to estimate the single-sample networks. Therefore, it is important to consider what samples are best to include in this background dataset. The background dataset can exist of a heterogeneous set of samples—for example, by including data derived from multiple conditions, tissues, or time points—or a homogeneous set of samples, such as multiple samples from the same specific condition.

We previously used simulated data to compare LIONESS networks built on heterogeneous or subtype-specific background data, including subtypes that made up different proportions of the data (Kuijjer et al., 2019a). We evaluated the accuracy of capturing edges in the gold standard, both when modeling LIONESS networks on the complete, or subtype-specific background. We found minimal differences in accuracy. Sometimes LIONESS networks modeled on the complete background performed better and other times networks modeled on the subtype-specific background performed better.

It is important to keep in mind that the decision on whether to use a heterogeneous or homogeneous dataset for the background depends on what types of edges are most relevant for your scientific question. For example, including only samples from a specific subtype will allow you to better investigate subtle differences in edges within that population of samples. However, if you include a heterogeneous set of samples in the background, it will also be possible to distinguish edges in that subtype from edges in the remaining samples (see also Figure S4C in Kuijjer et al., 2019a).

How LIONESS performs with different types of backgrounds also depends on the underlying network reconstruction algorithm used with LIONESS. Many network reconstruction algorithms require at least some heterogeneity across samples in a dataset to estimate edges. Thus, using a heterogeneous background may lead to more accurate edge predictions compared to using a homogeneous background; the latter may not have enough variability for the network reconstruction algorithm to accurately estimate edges.

### Preprocessing the Input Data

One of the most challenging steps in running a network reconstruction approach is preprocessing the data correctly to facilitate downstream interpretation and analysis. Before running LIONESS, make sure that all of the input data is in the correct format for the selected network reconstruction approach and that the required computational resources are available (see the *Running LIONESS* section below).

#### Filtering and Aligning the input data

The input data may need to be filtered before running LIONESS. This again depends on the chosen aggregate network reconstruction algorithm, the type of input data that is being used to model the networks, and the number and type of edges that are needed for downstream analysis. Filtering will also reduce the size of the LIONESS networks, so that less computational resources and storage space are required. However, filtering can also remove signal that may be critical when applying genome-wide network approaches and downstream pathway analysis. We describe some of these choices for Pearson and PANDA networks below.

##### Pearson correlation

When applying LIONESS to Pearson correlation networks, the single-sample networks can get very large, especially when modeling genome-wide networks. To illustrate this, a Pearson correlation network between 20,000 protein-coding genes would result in 20,000 choose 2 gene pairs, which is nearly 200 million edges. In the example here, we selected the 1000 most variable genes, based on their standard deviation across all samples, before applying LIONESS. An alternative approach could be to select L1000 genes (Subramanian et al., 2017), or genes from a specific biological pathway of interest. Selecting the top 1000 variable genes results in a network consisting of 1000 choose 2 edges, which is equal to 499,500 edges. While this is still a large number of edges, it is considerably fewer than a genome-wide Pearson correlation network and will be easier to load into memory and analyze.

##### PANDA

Applying LIONESS to PANDA networks involves a few additional steps, since PANDA matches targeting patterns in a prior regulatory network with gene co-expression and protein-protein relationships.

First, a prior regulatory network of putative (TF to gene) interactions needs to be defined. These can be obtained by scanning the promoter regions of genes for TF motifs from databases such as JASPAR (Fornes et al., 2019) or CISBP (Weirauch et al., 2014) using tools such as FIMO (Grant et al., 2011). Alternatively, information on TF binding to gene regulatory regions can be derived from ChIP-Seq data. Motif scans (or ChIP-Seq data) can be filtered, for example by thresholding based on p-value, or by defining a specific genomic region of interest. In the human example we show here, we downloaded prior regulatory networks from https://sites.google.com/a/channing.harvard.edu/kimberlyglass/tools/resources (“Human Motif Mappings” based on gene symbols, filename MotifPriorRefSeq_-1000_500_1e-3). Motif scans can be filtered, for example by thresholding based on p-value, or by defining a specific genomic region of interest. This motif scan was pre-filtered by selecting and discretizing matches of TF motifs from CISBP with p-values smaller than 1e-5 to a region of [-750, 250] around the gene’s transcription start site (TSS). The yeast prior regulatory network was processed as described in Kuijjer et al., 2019a.

Second, expression data is used by PANDA to estimate co-expression relationships. As lowly expressed and lowly variable genes help define the distributions of the co-expression values, it is generally advisable to include most genes in the input expression data. In our example PANDA networks, we therefore only filtered out genes that had zero counts in more than half of all samples.

Finally, an optional data type in PANDA is information on PPIs between TFs. This information can be obtained from publicly available databases, such as BioGrid (Oughtred et al., 2019) or StringDb (Szklarczyk et al., 2015), or from experimental evidence such as yeast-2-hybrid screens (Luck et al., 2020). Here, we used scores representing strengths of functional interactions from StringDb as the PPI input in PANDA.

Finally, when running PANDA it is useful to ensure that all of the input data is aligned. This is because PANDA uses matrix operations on the three input matrices. These matrices need to have the correct dimensions, with the same set of TFs being present in the PPI and prior regulatory network, and the same set of target genes in the prior regulatory network and the expression data. Different implementations of PANDA will do this alignment automatically, but we often suggest additional filtering steps to better understand the implications for network reconstruction and analysis. In the example we show here we filtered genes in the expression data that were not targeted by any TF in the prior regulatory network as well as genes in the prior network without expression data. Similarly, we only kept TFs in the PPI that were in the prior regulatory network. This is to ensure that, for each TF and each target gene, prior information on potential regulation is available to prime the message passing algorithm in PANDA. We note that PANDA models the role of TFs and genes separately, therefore the identifiers used for the TFs do not need to match those used for the target genes.

##### Tip

To make sure the same sets of TFs and target genes are being used in PANDA, it is useful to separately process each data type, and then perform an intersection of: (1) the target genes present in the prior regulatory network and the expression data, and (2) the TFs present in the PPI and prior regulatory network.

##### Tip

Filtering out specific genes, as well as filtering out edges in the prior regulatory network based on the promoter range and p-value cut-off may change the structure of the prior regulatory network, impacting the predicted aggregate PANDA network, and thus the LIONESS networks. If you are unsure how different filtering steps may influence the network, consider modeling and comparing aggregate networks built with different filtering steps before applying LIONESS.

#### Preparing the data for LIONESS

It is important to preprocess the input data before running an aggregate network model. For transcriptomic data, this may involve batch correction, normalization, and log-transformation or other variance stabilizing normalization methods. As with data filtering, how the data is preprocessed may affect the aggregate network, and therefore also the LIONESS networks inferred from the aggregate network model.

##### Pearson correlation

As with general gene expression analysis, the input expression data for co-expression network modeling should be log-transformed to reduce skewness in the data. The data should also be normalized so that measurements are comparable across samples, and, if possible, batch-corrected to reduce technical variability. Several normalization and transformation methods exist that are specifically designed to normalize and reduce variance-mean relationships in microarray and RNA-Seq data. In principle, any method that corrects for technical artifacts in the data and that makes expression data comparable across samples can be used. However, keep in mind that the technique of normalization selected could have an effect on the output co-expression values (Hsieh et al., 2021). It is therefore important to evaluate the effect the chosen preprocessing technique may have on the chosen network reconstruction algorithm to be used with LIONESS.

##### PANDA

When modeling LIONESS networks with PANDA as the selected aggregate network reconstruction algorithm, the prior regulatory network and PPI data are generally “fixed” input datasets—they are the same for each sample—while the expression data is what distinguishes each individual sample from the other samples in the dataset. It is therefore important to preprocess the expression data as described above. As the distribution of the prior regulatory network will influence the final weight distribution in the network modeled with PANDA, it is important to think about whether the scores in the prior regulatory network should be normalized as well. For example, when using data derived from ChIP-Seq experiments, normalization before peak calling may affect the prior regulatory network. Moreover, by default, PANDA assigns a score of 1 to self-interactions in the PPI. As the range of the PPI scores in StringDb is [0,1000], we generally convert these to values between [0,1] by dividing the scores by 1000 (Sonawane et al., 2017).

##### Tip

To understand how the network reconstruction approach you select may be affected by data preprocessing, it can be helpful to directly compare the aggregate network model inferred with and without different preprocessing steps. We recommend doing this step-by-step, so that you can get an understanding of how each step influences the aggregate network.

#### Dealing with sparsity and/or missing values in the input data

Many network reconstruction methods rely on the variability across samples in a dataset to infer interactions between genes. Applying such aggregate network reconstruction algorithms to sparse data or data with missing values can therefore return NAs in the output aggregate network, which may in turn affect the networks returned by LIONESS. This can happen when modeling, for example, genome-wide networks based on RNA-Seq or single-cell RNA-Seq data, as these data often include genes with tied counts. This is generally avoided by filtering out genes with zero counts in a certain percentage of samples in the dataset (see also the section *Filtering the input data* above). When filtering out genes, it is important to remember that there needs to be enough variability across samples after each sample is removed from the background dataset. For example, when applying LIONESS to model Pearson correlation networks based on RNA-Seq data, you need to make sure that each gene pair has at least four matching samples with different real values, as Pearson correlation requires three samples with variable expression. If such instances cannot be avoided, any missing values (NAs) in the aggregate network can be substituted with a value that indicates no co-expression/regulation (Lopes-Ramos et al., 2020) (e.g. a Pearson correlation coefficient of 0).

In addition to NAs that might arise in the aggregate network due to sparsity of the input data, the input data itself can include NAs, which might result in issues when calculating the aggregate network if the chosen reconstruction approach cannot handle missing information. There are different ways to deal with NAs in data, including adding a pseudocount before log transformation of RNA-Seq count data, imputing missing values, or excluding samples with NAs on an edge-by-edge basis during network modeling; in our analyses we generally choose to use a pseudocount so that missing values are converted to zeros after log transformation. How to deal with NAs depends on whether you believe values are missing due to biological reasons, or whether they are undetected due to technical bias or the stochastic nature of sampling (Silverman et al., 2020). While adding a pseudocount to RNA-Seq count data is standard practice, other methods have not yet been tested in the context of LIONESS, and we would recommend using them with caution.

##### Tip

Missing values (NAs) in the aggregate networks used by LIONESS may be unavoidable. For example, if a gene has the same expression value across all samples, the standard deviation is zero and correlation cannot be computed (see the section *Dealing with sparsity and/or missing values in the input data* for more information). One way to deal with this in Pearson correlation networks is to impose a value of 1 for diagonal elements, and zero for missing values.

### Running LIONESS

Once all the input data is ready, you are ready to run LIONESS. First, you will need to calculate an aggregate network reconstruction approach using all samples, then remove one sample at a time, rerun the reconstruction approach on all samples except for the one removed, and apply the LIONESS equation. While this can be easily coded within another computational pipeline, LIONESS is also available in several programming languages, including R (Kuijjer et al., 2019b), Python (van IJzendoorn et al., 2016), and MATLAB (see also netzoo.github.io/), for use with Pearson correlation and PANDA. Note that different implementations of the same network reconstruction algorithm often have various trade-offs. For example, the MATLAB implementation of PANDA, which we use here, is memory intensive, but faster than implementations of the algorithm in other programming languages (Glass et al., 2015). Tutorials with examples on how to run the method are available in the vignettes of each package, as well as in NetZoo (netzoo.github.io/).

#### Computational load

LIONESS’ computational load depends on the computational load of the chosen network reconstruction algorithm. LIONESS requires running the chosen method (*N* + 1) times – once to model an aggregate network, as well as *N* times to model networks with each sample iteratively removed. Therefore, the total run-time of LIONESS can be approximated by calculating the run-time of the aggregate network and multiplying this by *N* + 1. However, the exact run-time will depend on the precise implementation of LIONESS.

Running times to read in the data, run the aggregate network model, and print out the network, on one core, on a server with 128 Intel Haswell cores, 1Tb of RAM:

##### Tip

Before running LIONESS, estimate the running time and memory that is needed for modeling and printing an aggregate network with your network reconstruction algorithm of choice. As LIONESS will repeat this *N* + 1 times, you can get an estimate of how long it will take to run and print all of the single-sample networks.

##### Tip

Before estimating single-sample networks for all samples in an input dataset, it is useful to apply LIONESS to only a few (3-10) samples. This will help identify and track down issues related to the interface between the input data, the selected network reconstruction approach, and the LIONESS procedure with significantly less time and resources than if it were first applied to all samples.

##### Tip

If the resources are available, consider parallelizing the code. For example, modeling the human aggregate PANDA network in MATLAB took only 436s (7m16s), 289s (4m49s), 205s (3m24s), and 145s (2m25s) when using 2, 4, 8, or 16 cores at the same time, respectively.

#### Printing and storing the networks

Depending on the computing resources available, the network reconstruction algorithm of choice, the size of the network, and the number of samples in the expression dataset, running LIONESS may take from several minutes to several days or even weeks. When working with a smaller dataset, it may be easiest to first calculate all the LIONESS networks and then print them to a single large file. However, when working with large sample sizes and/or large networks, it may be best to print out each single-sample network individually. This makes it easier to stop and resume the network modeling when the computation time is long and may be helpful in case the computing system that is used to model the networks experiences any issues. In addition, if the size of the network being estimated is large, then holding it in memory may use up valuable computational resources. Printing each single-sample network to its own file will slightly increase the run-time of LIONESS but will also free that memory to be used for other tasks. See for example the difference in peak memory usage between running just the aggregate network compared to all of the LIONESS networks using lionessR (Table 1). In some cases, it may also be useful to only print a portion of the network for downstream analysis (for example, edges with a weight above a threshold in the aggregate model built using all samples, or edges connected to genes of interest).

**Table 1.**
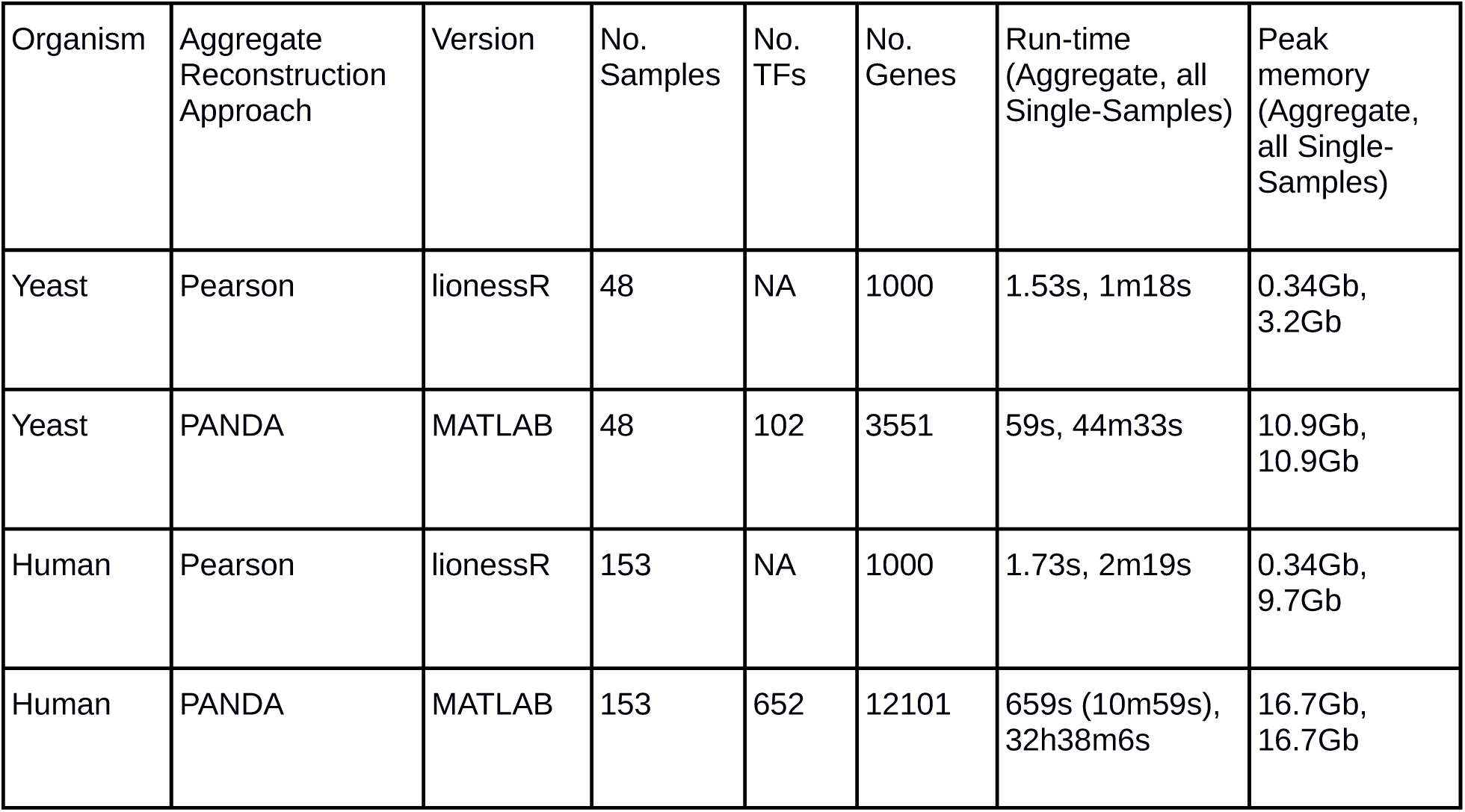
Running times and maximum amount of memory used for the network reconstruction approaches and data network reconstruction approaches and data discussed in this manuscript. Note the runtimes of LIONESS are system-and implementation-specific.

Printing the networks separately may make downstream parsing of the data a bit more challenging in some respects and less challenging in others. For example, any comparisons across samples / networks will require querying multiple files. It is possible that printing to separate files will take up more space on the disk, especially if the edge information (e.g. which genes, transcription factors, or other molecules are connected) is repeated in every file. However, it may also make data integrity to be harder to maintain, as it risks edges being difficult to interpret if the edge information is only printed to a single “master” file. In the latter case additional care will be critical when performing downstream analysis to ensure the information is correctly realigned. On the other hand, having each network in a separate file may help avoid memory issues in downstream analysis of the networks, since the entire set of networks do not have to be simultaneously read into memory.

### Expected outcomes

#### Initial assessment of the aggregate network

Before assessing the single-sample networks constructed by applying LIONESS, it is important to gain familiarity with features of the aggregate network obtained when applying the chosen network reconstruction approach (Step 1 of the procedure). Performing a baseline assessment of the aggregate network (for example, looking at the distribution of its edge weights) can help identify potential challenges that should be solved before applying LIONESS, such as missing weight assignments for certain edges, or an error in the computation of the aggregate network that needs to be corrected before using it iteratively with LIONESS.

#### Assessment of the single-sample networks

Before applying more complex analyses, there are several plots we generally recommend making in order to gain a high-level understanding of the structure of the aggregate and single-sample networks. First, we recommend looking at the distribution of the edge weights across the networks. Many network reconstruction approaches estimate a score for all possible edges in a network. These scores often follow a distribution that will be reflected in the LIONESS-derived single-sample networks. For example, the weights of edges in a co-expression network are generally centered around zero and bounded by zero and one. In contrast, PANDA edge weight distributions are often bimodal, with the two peaks representing the sets of edges that either have, or do not have, evidence in the prior regulatory network. In this section, we describe and present several visualizations of the single-sample networks we obtained by applying LIONESS to model Pearson correlation and PANDA networks on yeast data, as described above.

Figure 2A shows the distribution of edge weights in the aggregate co-expression network, the distribution of edge weights across all of the single-sample networks reconstructed using LIONESS, as well as the distribution of the weight of each edge when taking the mean across the single-sample networks. These distributions reveal important patterns in the LIONESS results. In particular, we can see that the single-sample networks retain a similar distribution as the aggregate network centered at zero. However, the edge weights of the single-sample networks are no longer bounded between -1 and 1. This may be important to consider in downstream analysis, for example when using tools typically applied to Pearson networks since these may assume boundedness.

**Figure 2.**
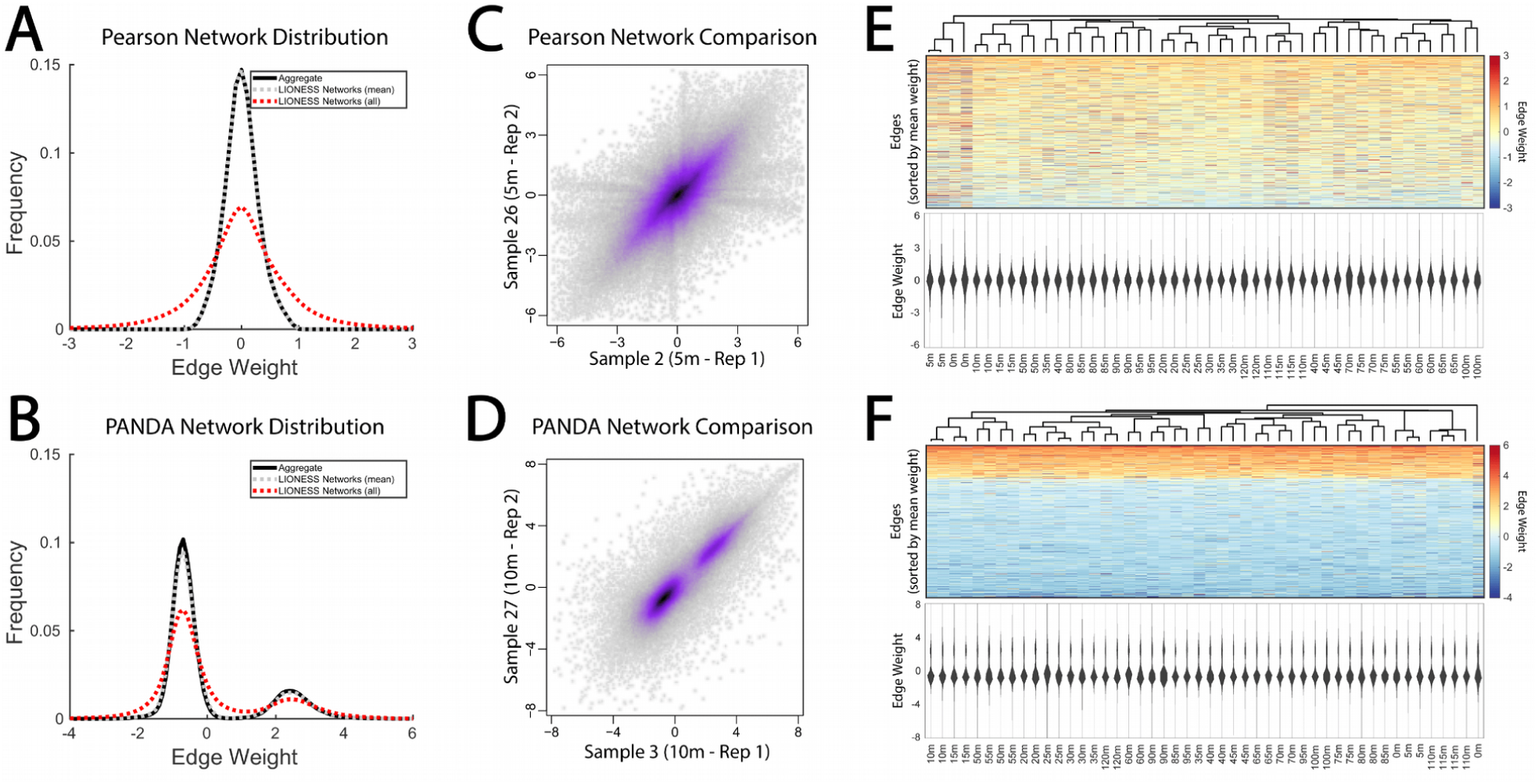
(A-B) Distribution of edge weights for aggregate and single-sample networks estimated using (A) Pearson correlation, and (B) PANDA. (C-D) Smooth scatter plots comparing two single-sample networks that correspond to technical replicates in the original expression data. This includes (C) single-sample networks based on Pearson correlation for the 5m time point, and (D) single-sample networks based on PANDA for the 10m time point. (E-F) Clustering and visualization of all the edges estimated by LIONESS when used with (E) Pearson correlation and (F) PANDA. In both cases, the heat map illustrates the weight of the estimated edges, with rows sorted by the average edge weight across all single-sample networks, and columns clustered using hierarchical clustering (Spearman distance, complete linkage). The violin plots show the distribution of edge weights in each of the single-sample networks.

Figure 2B shows these same distributions in the case of applying LIONESS to PANDA. Here we note that there are two classes of edges, one with significantly higher weight than the other. This is because PANDA uses prior information to upweight edges with evidence for TF binding. This information is propagated to the single-sample networks. Importantly, however, for both the Pearson and PANDA-derived networks the distribution of the mean-weights of edges across the single-sample networks is identical to that of the aggregate network. This is consistent with the underlying model used by LIONESS, which stipulates that a linear combination (average) of single-samples network models should be equal to an aggregate network model reconstructed using all samples (Kuijjer et al., 2019a).

Second, we recommend plotting the weights of edges in pairs of networks against each other. For example, Figures 2C-D shows networks reconstructed for the 5m and 10m time point between two technical replicates in the yeast data. Plotting pairs of networks in this way will provide a sense of how similar the single-sample networks are to one another. The single-sample networks derived using LIONESS include both the common structure that is shared across all the samples, as well as structure that is specific to that individual network. Thus, in most instances, comparing two single-sample networks will reveal a high level of similarity, with many edges—those that are not specific to either sample but common to both—falling along the diagonal. This type of visualization is also a useful and quick way to identify potential issues with the computation of the networks. For example, if a pair of single-sample networks are identical, then there may be an issue with correctly iterating through the individual samples when computing the networks.

Finally, given sufficient computational resources, it can also be beneficial to cluster and visualize the distribution of edges across all of the single-sample networks. Figures 2E-F each include a clustered heat map illustrating the weight of all edges (rows) across all the single-sample networks, as well as violin plots showing the distribution of those edge weights in each of the single-sample networks. Of note, several individual single-sample Pearson networks (Figure 2E) appear to have a more heterogeneous edge weight distribution compared to the other samples. Although this may be biologically relevant or informative of an underlying phenotype, it may also need to be considered in downstream analysis. Similarly, in the heat map showing the edge weights when applying LIONESS together with the PANDA network reconstruction approach (Figure 2F), the bimodal distribution of edges is striking. Although this is expected given the aggregate and single-sample network distributions in Figures A-B, it also reinforces the importance of understanding the structure of the network predicted by the aggregate reconstruction approach used with LIONESS, the dependence of the single-sample networks on the output of that approach, and taking that into consideration when performing downstream analyses.

Finally, we also recommend comparing the clustering of LIONESS networks to a clustering of the input data used to construct those networks (in these examples, the input gene expression data). These clusterings can sometimes be quite distinct, as we observed in our Methods paper (Kuijjer et al., 2019a) when analyzing lymphoblastoid networks constructed using LIONESS. In this case differences in the targeting patterns of genes in the networks between clusters was highly associated with relevant biological function, whereas differences in the expression levels of genes between clusters identified using the input gene expression were not.

### Downstream analysis

#### Evaluating and comparing single-sample networks

After modeling and performing initial assessments of the LIONESS networks, you can start downstream analysis. There are many different approaches that can be used to analyze single-sample networks—the best approach to use often depends on what question you would like to answer. For example, similar to differential expression analysis, you can compare networks belonging to two or more groups of samples to determine if there are any statistical differences between these groups. Such analyses can be performed on different network metrics characterizing information about the edges or nodes. This includes, but is not limited to, measures such as the edge weight (Sonawane et al., 2017), and node targeting (Weighill et al., 2021).

One advantage of determining differences in edge weights is that it can help identify specific interactions that are different between sample groups. However, this analysis also may involve testing differences for a very large number of edges. Thus, the number of false positive findings may be large when thresholding based on p-value and correction for multiple testing is necessary (Noble, 2009). In order to alleviate the multiple testing burden, it might be useful to reduce the number of edges prior to running a statistical test. For example, if there are specific genes or pathways you would like to focus on, you could restrict your analysis to the edges connected to those genes or pathways. When analyzing regulatory networks, you can also reduce the number of edges by focusing on putative interactions—interactions that are supported by potential TF binding in the promoter of the target gene—in the statistical comparison. While this does not allow for the identification of potential new binding events, it will likely significantly reduce the number of tests performed.

Another approach to determine differences in networks based on comparing edge weights is to separately analyze all of the edges that are connected to a specific node. For example, we previously ran Gene Set Enrichment Analysis (GSEA) on differential “targeting patterns” (differential edge weights connected to a TF) to characterize tissue-specific regulation of biological pathways by each individual TF (Sonawane et al., 2017). We point out, however, this type of analysis may require large computational resources. Preselecting the TFs, for example based on if changes in their targeting is correlated with changes in downstream gene expression (Lopes-Ramos et al., 2020), can help reduce the computational burden.

Networks can also be compared using node metrics. A widely used metric is the degree, or the sum of all edge weights connected to a node. For bipartite transcriptional networks modeled with PANDA, both the TF’s out-degree and the target gene’s in-degree (also called the Gene Targeting Score) can be calculated (Weighill et al., 2021). These measures are powerful, as they are relatively quick to calculate. They also allow one to analyze the networks at the gene level and perform tests such as Gene Set Enrichment Analysis to identify differentially regulated pathways. Other node metrics that can be used include node betweenness, closeness, or eigenvalue centrality. For example, we previously analyzed gene targeting as well as gene betweenness in PANDA regulatory networks and found that these metrics were associated with the gene’s tissue-specific expression (Sonawane et al., 2017). While the degree is often quick to calculate, other metrics can be more computationally challenging to run. For example, to calculate the betweenness, an algorithm will need to calculate the number of shortest paths that run through each node. Thus, while it is feasible to apply these node metrics to relatively small networks, they can be challenging to calculate for large, weighted, or complete graphs— characteristics that often describe the networks estimated by LIONESS. In addition, calculating these metrics may require more computational resources or longer run-times, issues that will be exacerbated when calculating them for a large number of single-sample networks.

Different statistical tests can be used to compare edge weights and node metrics between groups of single-sample networks. The selected test should be appropriate for the type of single-sample networks that are being compared and will depend on the distribution of scores obtained from applying LIONESS to a particular network reconstruction model and data type(s). It is important to note that, due to the nature of the LIONESS equation, there could be outliers in the distribution of edge weights that may violate assumptions of certain statistical tests, for example those that require a normal distribution (see the section *Outliers in single-sample networks* below). On occasion, the LIONESS equation may result in edge weights that are not normally distributed across samples, in which case it may be more appropriate to use nonparametric statistics when comparing single-sample edge weights. However, in our experience, node degree is often normally distributed across samples and can generally be compared with, for example, Student’s t-statistic or linear regression-based approaches (Lopes-Ramos et al., 2018, 2020, 2021).

##### Tip

Visualize the distributions of network metrics you would like to compare, so that you can select a statistical test whose underlying assumptions fit the data.

##### Tip

Before applying network metrics to individual single-sample networks, it is important to understand the expectations of the metric. For example, many network metrics only work for positive edge weights, but LIONESS networks might include negative edges. This is especially important to consider when using aggregate approaches that estimate both positive and negative edge weights (see Figure 2). If LIONESS networks include negative edges, a transformation of edge weights could be used, such as that applied in Sonawane et al., 2017.

#### Clustering single-sample networks

Networks modeled for single samples can be analyzed by applying clustering approaches to network edge weights, node degrees, or other network metrics. Clustering can help identify whether networks modeled on biological or technical replicates are reproducible (as we show in Figures 2E-F) or help find potential new groupings of samples (as in Kuijjer et al., 2019a). Multiple clustering algorithms, including hierarchical clustering, k-means clustering, non-negative matrix factorization (NMF), as well as non-parametric data visualization approaches such as t-distributed Stochastic Neighbor Embedding (t-SNE, van der Maaten and Hinton, 2008) and Uniform Manifold Approximation and Projection (UMAP, McInnes et al., 2018) could, in principle, be applied to single-sample networks.

Hierarchical clustering is relatively quick, which is an advantage when clustering multiple large networks. In addition, the number of clusters does not need to be defined beforehand. Hierarchical clustering measures the dissimilarity between samples. We recommend first determining which distance metric is most appropriate for your analysis. This will depend on the network reconstruction algorithm used with LIONESS, as well as on the specific network metric being clustered. Because LIONESS networks may include outliers (as we describe in the section *Outliers in single-sample networks*), ranked-based metrics such as Spearman distance are likely to better fit the underlying data than parametric approaches such as Pearson distance.

Methods such as k-means clustering and NMF require predefining the number of clusters (*k*). As this number is generally unknown, these methods are often run across a range of *k* values. The best clustering is then determined using various metrics, such as the cophenetic correlation or the Silhouette value. Because these methods generally need to be run over a range of *k* they can be slow. Therefore, they may not be the best approaches to use when clustering very large networks. Another thing to keep in mind is that NMF requires all input data to be non-negative. Therefore, depending on the aggregate reconstruction approach used, you may need to transform the LIONESS output to ensure positive edge weights (Sonawane et al., 2017).

Non-parametric methods such as tSNE and UMAP are designed to work well for complex, non-linear data and could therefore be powerful when visualizing sample dependencies in networks. However, we have not yet systematically evaluated them in the context of analyzing LIONESS networks.

#### Outliers in single-sample networks

We often observe a large range of edge weights across the different sample networks predicted by LIONESS, with the edges in a handful of single-sample networks having more extreme edge weights compared to the others. Quite often these edges are associated with a single-sample network that has a distinct distribution from the other single-sample networks (e.g. the network at the 0m time point in Figure 2F). Investigating the input data used by LIONESS to compute these “outlier” edges typically reveals a strong sample-specific difference. For example, if most samples indicate a strong positive correlation between a pair of genes, but one sample bucks this trend, the edge between the pair of genes in that single-sample Pearson network will be negative while the weight of the edge in the rest of the networks will be centered around one. However, even though they correctly reflect the underlying data, these edges can also dominate clustering results, potentially leading to a poor grouping of the samples. To address this issue, one option is to quantile normalize edge weights or network metrics to standardize across the single-sample networks (Lopes-Ramos et al., 2018). Alternatively, as we did in Sonawane et al., 2017, you could compare edges and other network metrics using nonparametric instead of parametric tests.

#### Other types of network analyses

Supervised statistical comparisons of edge or node metrics, or unsupervised analyses through clustering of LIONESS networks, are good first steps for identifying biological differences between single-sample networks. However, a multitude of network analysis methods exist that could also potentially be applied to LIONESS networks. For example, the topology of each of the single-sample networks could be analyzed using community structure detection methods to identify modules of strongly connected nodes. These modules may be enriched for specific biological properties (Platig et al., 2016; Fagny et al., 2017, 2020). In addition, community structure comparison methods (Padi and Quackenbush, 2018; Lim et al., 2020) can be used to identify differences between pairs of single-sample networks. Although these methods are generally designed to compare two networks, they could be used to iteratively compare the community structure of each of the single-sample networks with that of the aggregate network. Other potential extensions of LIONESS network analysis include associating network structure with sample-specific node information (such as gene expression levels), layering in multiple types of edges, using the networks in machine learning approaches to build predictors (e.g. of clinical outcomes) or classifiers, applying network fusion approaches, or using network propagation approaches (Cowen et al., 2017). For more information and tips on different network analysis approaches, we would like to refer the reader to Miele et al., 2019.

## Discussion

LIONESS is a versatile method that can be combined with different types of aggregate network reconstruction approaches to model networks for individual samples. In this work, we demonstrate how to apply LIONESS to co-expression and gene regulatory networks in human and yeast and provide examples of how to analyze the single-sample networks predicted by LIONESS. We also discuss how the chosen network reconstruction algorithm, input data, and data preprocessing steps might influence the single-sample network estimates, as well as potential computational challenges that might arise when modeling and analyzing networks for individual samples. Because LIONESS can be applied to very different types of network reconstruction algorithms and data, we recommend that the user explores how the data and data preprocessing may affect the output networks, as well as identify the downstream network analysis approach that is most appropriate to answer their question of interest. These decisions are critical to gaining important biological insights, as we and others have in our previous applications of the method (Kuijjer et al., 2019a; Sonawane et al., 2019; Fagny et al., 2020; Lopes-Ramos et al., 2018, 2020, 2021, Pham et al., 2021).

## Acknowledgments

MLK is supported by the Norwegian Research Council, Helse Sør-Øst, and University of Oslo through the Centre for Molecular Medicine Norway (187615), as well as the Norwegian Research Council (313932) and the Norwegian Cancer Society (214871). KG is supported by the US National Heart, Lung, and Blood Institute of the National Institutes of Health (K25HL133599).

## Author Contributions

Conceptualization, Methodology, Software, Formal Analysis, Resources, Writing—Original Draft, Writing—Review & Editing, Visualization, Funding Acquisition: MLK and KG.

